# Concurrent duplication of the Cid and Cenp-C genes in the Drosophila subgenus with signatures of subfunctionalization and male germline-biased expression

**DOI:** 10.1101/134817

**Authors:** José R. Teixeira, Guilherme B. Dias, Marta Svartman, Alfredo Ruiz, Gustavo C. S. Kuhn

**Affiliations:** Departamento de Biologia Geral, Universidade Federal de Minas Gerais, Belo Horizonte, Minas Gerais, Brazil, Postal Code: 31270-901; Departament de Genètica i de Microbiologia, Universitat Autònoma de Barcelona, Bellaterra (Barcelona), Spain, Postal Code: 08193

**Keywords:** CenH3, Cenp-C, gene duplication, centromere, kinetochore, Drosophila

## Abstract

Despite their essential role in the process of chromosome segregation in eukaryotes, kinetochore proteins are highly diverse across species, being lost, duplicated, created, or diversified during evolution. Based on comparative genomics, the duplication of the inner kinetochore proteins CenH3 and Cenp-C, which are interdependent in their roles of stablishing centromere identity and function, can be said to be rare in animals. Surprisingly, the *Drosophila CenH3* homolog *Cid* underwent four independent duplication events during evolution. Particularly interesting are the highly diverged and subfunctionalized *Cid1* and *Cid5* paralogs of the *Drosophila* subgenus, which show that over one thousand *Drosophila* species may encode two *Cid* genes, making those with a single copy a minority. Given that CenH3 and Cenp-C likely co-evolve as a functional unit, we investigated the molecular evolution of *Cenp-C* in species of *Drosophila*. We report yet another *Cid* duplication within the *Drosophila* subgenus and show that not only *Cid*, but also *Cenp-C* is duplicated in the entire subgenus. The *Cenp-C* paralogs, which we named *Cenp-C1* and *Cenp-C2*, are highly divergent. The retention of key motifs involved in centromere localization and function by both Cenp-C1 and Cenp-C2 makes neofunctionalization unlikely. In contrast, the alternate conservation of some functional motifs between the proteins is indicative of subfunctionalization. Interestingly, both *Cid5* and *Cenp-C2* are male germline-biased and evolved adaptively. Our findings point towards a specific inner kinetochore composition in a specific context (i.e., spermatogenesis), which could prove valuable for the understanding of how the extensive kinetochore diversity is related to essential cellular functions.

## Introduction

During eukaryotic cell division, accurate chromosome segregation requires the interaction of chromosomes with the microtubules from the spindle apparatus. This interaction is mediated by the kinetochore, a multiprotein structure that is hierarchically assembled onto centromeres. Upstream in the assembly of the kinetochore are CenH3 and Cenp-C, two interdependent proteins in their roles of establishing centromere identity and function. CenH3 is the histone H3 variant found in centromeric nucleosomes and, therefore, considered the centromere epigenetic marker (Dalal *et al*. 2007). During kinetochore assembly, Cenp-C binds to CenH3 and recruits other kinetochore proteins (Przewloka *et al*. 2011; Liu *et al*. 2016). CenH3 and Cenp-C are fundamentally interdependent because the centromeric localization of one depends on the centromeric localization of the other (Erhardt *et al*. 2008; Orr and Sunkel 2011). This interdependence is also illustrated by the fact that both CenH3 and Cenp-C have similar phylogenetic profiles (i.e., they have similar patterns of presence and absence across the eukaryotic evolutionary tree) and likely co-evolve as a functional unit (van Hooff *et al*. 2017). One interesting case is that seen in insects, where CenH3 was lost independently five times, and in all these cases Cenp-C was also lost (Drinnenberg *et al*. 2014).

Despite the essentiality of centromeres, both centromeric DNA (CenDNA) and proteins are remarkably diverse (Henikoff *et al*. 2000; Talbert *et al*. 2004; Plohl *et al*. 2008). This rapid evolution despite the expectation of constraint is referred to as the “centromere paradox” (Henikoff *et al*. 2001). This paradox may be explained by the centromere drive hypothesis, which proposes that genetic conflicts during female meiosis drive centromere evolution (Henikoff *et al*. 2001; Dawe and Henikoff 2006).

In the female meiosis of animals and plants, the meiotic spindle fibers are asymmetric in a way that one pole will originate a polar body and the other will give rise to the oocyte. As a result, there is potential for non-mendelian (biased) inheritance if a pair of homologous chromosomes have kinetochores that interact unequally with the spindle fibers (Ross and Malik 2014). The heterogeneity in kinetochore function between homologs is a result of differences in abundance of centromeric DNA sequences. One homolog may have a ‘strong’ centromere, which has an expanded cenDNA that recruits more kinetochore proteins and delivers its chromosome into the oocyte at > 50% frequency, or a ‘weak’ centromere, which has a contracted cenDNA that in turn recruits less kinetochore proteins and delivers its chromosome into the oocyte at < 50% frequency (Iwata-Otsubo *et al*. 2017). However, the spread of expanding centromeres throughout a population might be accompanied by deleterious effects, such as increased male sterility or a skewed sex ratio (Fishman and Saunders 2008; Rutkowska and Badyaev 2008; Malik and Henikoff 2009). The centromere drive hypothesis proposes that changes in CenH3 and Cenp-C related to more ‘flexible’ DNA-binding preferences are expected to counteract the transmission advantage gained by expanded centromeres and diminish the associated deleterious effects, thus restoring meiotic parity for both homologs (Henikoff et al. 2001; Dawe and Henikoff 2006).

The kinetochore is highly diverse across species, with proteins being lost, duplicated, created, or diversified during evolution (van Hooff *et al*. 2017). Given that data directly supporting a correlation between the evolution of cenDNA, CenH3 and Cenp-C are still absent, it is not known if and how such structural divergence is related to centromere drive suppression. However, the subfunctionalization of *CenH3* paralogs in some lineages of *Drosophila* has been hypothesized to be linked to centromere drive suppression. Kursel and Malik (2017) have recently reported that the *Drosophila CenH3* homolog *Cid* underwent four independent duplication events during evolution, and some *Cid* paralogs are primarily expressed in the male germline and evolve under positive selection (Kursel and Malik 2017). These duplications could have allowed the rapid evolution of centromeric proteins without compromising their essential function by separating functions with divergent fitness optima. The existence of germline-biased *CenH3* duplicates (which do not interfere with essential mitotic functions) in genetically tractable organisms provides an opportunity to study the functional consequences of the genetic variation for kinetochore-related processes.

Given the interdependence between CenH3 and Cenp-C, we decided to further analyze the molecular evolution of the *Cid* and *Cenp-C* genes in *Drosophila* species. Here, we report a novel *Cid* duplication within the *Drosophila* subgenus and show that not only *Cid*, but also *Cenp-C* is duplicated in the entire *Drosophila* subgenus. The *Cid* and *Cenp-C* paralogs likely subfunctionalized, as some motifs are alternatively conserved between the paralogs. Interestingly, both the *Cid* and *Cenp-C* duplications generated copies that are male-biased and evolve under positive selection. Our findings point towards a specific kinetochore composition in a specific context (i.e., the male germline), which could prove valuable for the understanding of how the extensive kinetochore diversity is related to essential cellular functions.

## Results and Discussion

### *Cid1* was replaced by a new paralog in a clade within the *Drosophila* subgenus

Duplicate *Cid* genes exist in *D. eugracilis* (*Cid1, Cid2*) and in the *D. montium* subgroup (*Cid1, Cid3, Cid4*), both within the *Sophophora* subgenus, and in the entire *Drosophila* subgenus (*Cid1, Cid5*). In all analyzed species from the *Drosophila* subgenus, *Cid1* is flanked by the *cbc* and *bbc* genes, and *Cid5* is flanked by the *Kr* and *CG6907* genes (Kursel and Malik 2017). As expected, we found two *Cid* genes while looking for the orthologs of *Cid1* and *Cid5* in the assembled genomes of two cactophilic species from the *Drosophila* subgenus, *D. buzzatii* and *D. seriema* (*repleta* group). Surprisingly, while one of the genes is present in the expected locus of *Cid5*, the other one is located in an entirely different locus, flanked by the *CG14341* and *IntS14* genes. We named this new paralog as *Cid6*.

By investigating the *Cid1* locus of *D. buzzatii*, we found a myriad of transposable elements (TEs) surrounding a 116-bp fragment of the original *Cid1* gene (fig. 1, upper panel). Due to fragmentary genome assembly, the *Cid1* locus of *D. seriema* could not be identified. Both *Cid5* and *Cid6* of *D. buzzatii* and *D. seriema* share ∼40% amino acid identity but, in contrast, *Cid6* of each species and *Cid1* of the closely related *D. mojavensis* are much more similar, sharing ∼80% identity. Fluorescent *in situ* hybridizations on polytene chromosomes showed that *Cid6* is distal (in relation to the chromocenter) in the Muller element B of *D. buzzatii* and *D. seriema*, and that *Cid1* is proximal in the Muller element C of *D. mojavensis* and the outgroup *D. virilis* (fig. 1, lower panel). Therefore, we inferred that *Cid1* was degenerated by several TE insertions after the origin of *Cid6* by an inter-chromosomal duplication of *Cid1* in the lineage that gave rise to *D. buzzatii* and *D. seriema*. The time of divergence between *D. buzzatii* and *D. seriema* has been estimated at ∼4.6 mya, and the divergence between them and the closely related *D. mojavensis* has been estimated at ∼11.3 mya (Oliveira *et al*. 2012). Therefore, the *Cid1* duplication that gave rise to *Cid6* happened between ∼4.6 and 11.3 mya.

**Figure 1.**
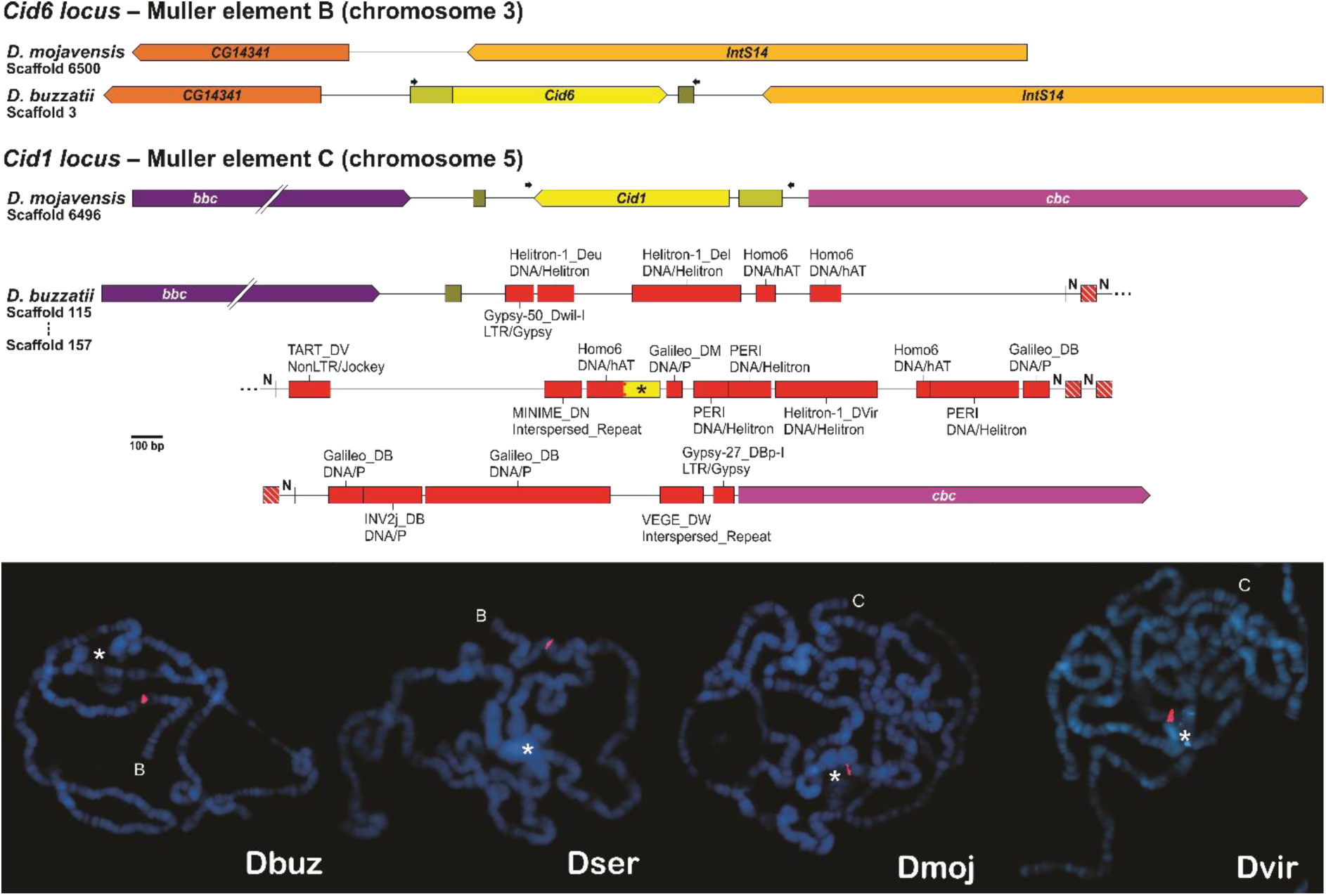
*Cid1* degenerated after the inter-chromosomal duplication event giving rise to *Cid6*. (Upper panel) Comparison between the *Cid1* and *Cid6* loci of *D. buzzatii* and the corresponding regions of *D. mojavensis*. The black asterisk indicates a fragment of *Cid1*, ‘N’ indicates unidentified nucleotides, red boxes indicate transposable elements, and arrows indicate primers used for the fluorescent *in situ* hybridization (FISH) experiments. (Lower panel) FISH on polytene chromosomes of *D. buzzatii* (Dbuz) and *D. seriema* (Dser) using *Cid6* probes, and of the closely related *D. mojavensis* (Dmoj) and the outgroup *D. virilis* (Dvir) using *Cid1* probes. The chromosome arm in which the *Cid* probe hybridized (red signal) is indicated by a letter representing the corresponding Muller element. The chromocenter, a region in which all centromeres bundle together, is indicated by a white asterisk. (Note: the chromocenter of *D. buzzatii* and *D. mojavensis* ruptured during the fixation step).

Why *Cid6* remained while *Cid1* degenerated? The *Cid1* locus of *D. buzzatii* is located in the most proximal region of the Muller element C (scaffold 115; Guillén *et al*. 2014), which is very close to the pericentromeric heterochromatin where TEs are highly abundant (Pimpinelli *et al*. 1995; Casals *et al*. 2005; Rius *et al*. 2016). Natural selection is known to be less effective in pericentromeric and adjacent regions due to low rates of crossing-over (Zhang and Kishino 2004; Clément *et al*. 2006; Comeron *et al*. 2012; Nambiar and Smith 2016). Thus, it is reasonable to suggest that the presence of an extra copy of *Cid1* (i.e., *Cid6*) in Muller element B alleviated the selective pressure on *Cid1* in Muller element C, whose proximity to the pericentromeric heterochromatin fostered its degradation by several posterior TE insertions.

### *Cenp-C* is duplicated in the *Drosophila* subgenus

It has been recently shown that the *Drosophila CenH3* homolog *Cid* underwent duplication events during evolution (Kursel and Malik, 2017). Given that CenH3 and Cenp-C are interdependent and coevolve as a functional unit, we investigated if *Cenp-C* was also duplicated in *Drosophila* species where *Cid* was duplicated.

In *D. eugracilis*, in species from the *montium* subgroup, and in all the other species of the *Sophophora* subgenus we found only one copy of *Cenp-C*, which is always flanked by the *5-HT2B* gene. On the other hand, in the species of the *Drosophila* subgenus we found two copies of *Cenp-C* with ∼52% nucleotide identity, which we named *Cenp-C1* and *Cenp-C2*: the former is flanked by the *5-HT2B* and *CG1427* genes, and the latter is flanked by the *CLS* and *RpL27* genes. A maximum likelihood tree showed that *Cenp-C* was likely duplicated after the split between the *Sophophora* and *Drosophila* subgenera but before the split between *D. busckii* and the other species of the *Drosophila* subgenus (fig. 2). Thus, we concluded that *Cenp-C2* originated from a duplication of *Cenp-C1* in the lineage that gave rise to species of the *Drosophila* subgenus, at least 50 mya (Russo *et al*. 2013).

**Figure 2.**
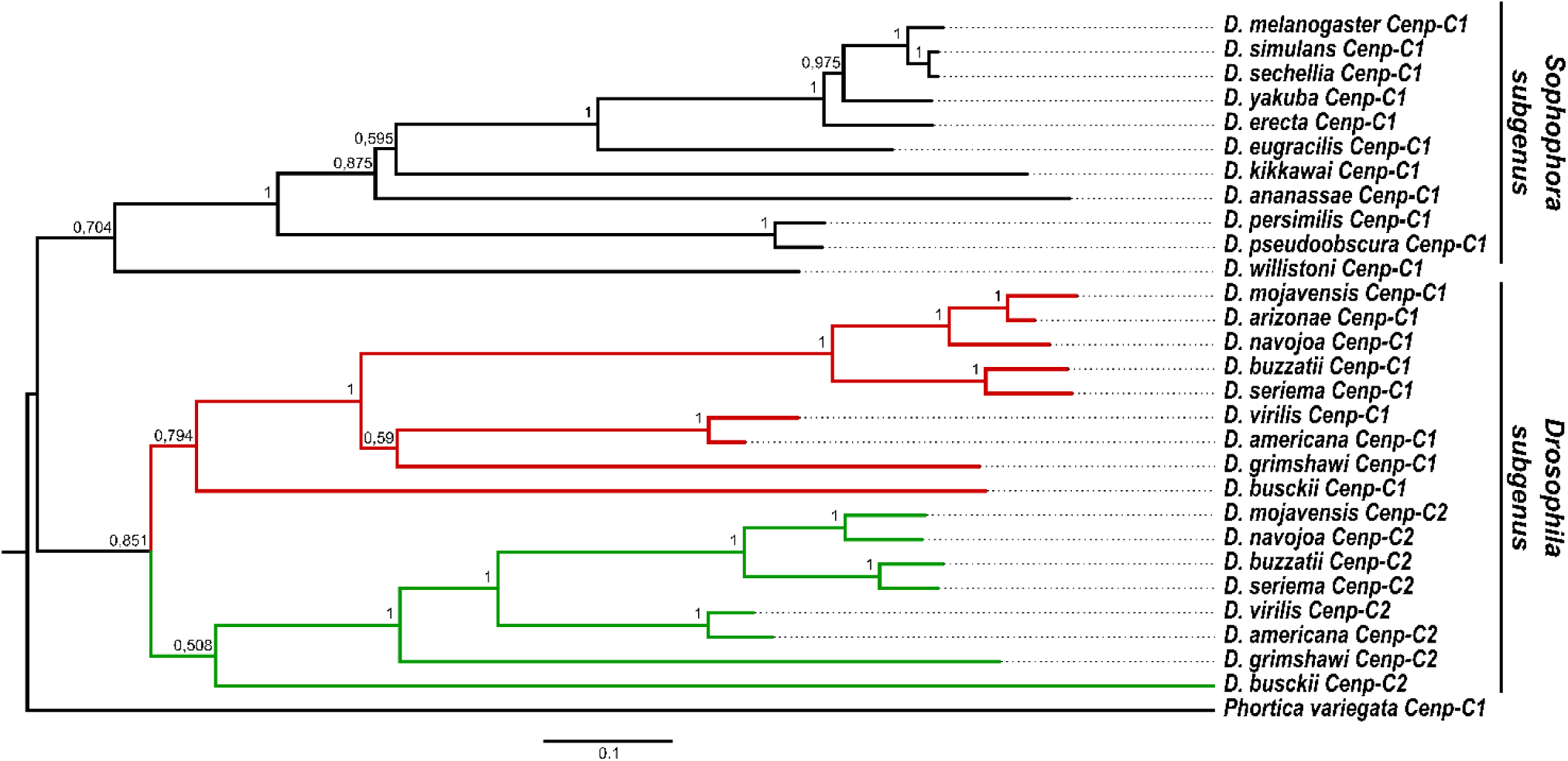
*Cenp-C1* duplicated in the lineage that gave rise to species of the *Drosophila* subgenus. Maximum likelihood tree of the *Cenp-C1* and *Cenp-C2* paralogs. Red and green branches respectively correspond to *Cenp-C1* and *Cenp-C2* sequences from species of the *Drosophila* subgenus. Bootstrap values are shown in each node. Scale bar represents number of substitutions per site.

Why *Cenp-C* is duplicated only in the *Drosophila* subgenus if *Cid* is also duplicated in *D. eugracilis* and in the *montium* subgroup? The fact that both *Cid* and *Cenp-C* duplicated in the *Drosophila* subgenus does not mean that there is a cause-and-effect relationship between the duplications. However, it probably means that the new paralogs influenced each other’s evolution. As a histone H3 variant, CenH3 has the C-terminal histone fold domain, which is reasonably conserved among species, and the N-terminal tail (NTT), which is highly variable among species (Henikoff *et al*. 2000). The NTT evolves in a modular manner, with four core motifs always conserved when there is only one Cid protein encoded in the genome (Kursel and Malik 2017). In *D. eugracilis*, the *Cid2* paralog functionally replaced the pseudogenized ancestral *Cid1* paralog. In species of the *montium* subgroup, these four motifs are alternated between the paralogs, which share ∼25% amino acid identity. In contrast, in species of the *Drosophila* subgenus, all four motifs are conserved in *Cid1* but only 1-2 are conserved in Cid5, with the paralogs sharing only ∼15% amino acid identity at their NTT. Therefore, we propose that if the NTT of Cid interacts with Cenp-C, a new *Cenp-C* copy would allow a higher divergence of the *Cid* paralogs by alleviating the selective pressure over the Cid/Cenp-C interaction, thus explaining the higher divergence of the *Cid1* and *Cid5* paralogs. However, future studies focusing on the specific interactions between Cid and Cenp-C shall shed light on the exact basis behind the flexibility of these two proteins during evolution.

### Some Cenp-C motifs are alternatively conserved between Cenp-C1 and Cenp-C2

Cenp-C was previously thought to be absent in *Drosophila* (Talbert *et al*. 2004), but it turned out that a protein that interacts with the regulatory subunits of separase is a highly divergent Cenp-C homolog (Heeger *et al*. 2005). The *D. melanogaster* Cenp-C1, as characterized by Heeger *et al*. (2005), has seven independent functional motifs, from N- to C-terminal: arginine-rich (R-rich), drosophilids Cenp-C homology (DH), AT hook 1 (AT1), nuclear localization signal (NLS), CenH3 binding (also known as the Cenp-C motif), AT hook 2 (AT2), and C-terminal dimerization (Cupin). The R-rich and DH motifs, as well as both AT1 and AT2 motifs (which may mediate binding to the minor grove of DNA), are functionally poorly characterized. However, all except AT1 appear to hold essential functions, as Cenp-C1 variants lacking these regions are unable to prevent phenotypic abnormalities in *Cenp-C1* mutant embryos (Heeger *et al*. 2005). In fact, it is known that the DH motif must be involved in the recruitment of kinetochore proteins (Przewloka *et al*. 2011; Liu *et al*. 2016). Furthermore, arginine 1101 (R1101), present in the CenH3 binding motif, is crucial for centromere localization (Heeger *et al*. 2005). Given the functional relevance of these motifs, we searched for them in both Cenp-C1 and Cenp-C2.

With the exception of *D. kikkawai* (from the *montium* subgroup), in which the AT2 motif is absent, all seven motifs are conserved in Cenp-C1 from all other species of the *Sophophora* subgenus. In contrast, the motifs are alternatively conserved between Cenp-C1 and Cenp-C2 in species from the *Drosophila* subgenus (fig. 3). Both Cenp-C1 and Cenp-C2 of all species have the DH, NLS, and CenH3 binding motifs (with the corresponding R1101 of *D. melanogaster*), but lack the AT1 motif. Furthermore, only Cenp-C2 has the R-rich and AT2 motifs conserved. Both Cenp-C1 and Cenp-C2 of most species have the Cupin motif, the exceptions being Cenp-C1 of *D. busckii*, which lacks the final half of it, and Cenp-C2 of *D. grimshawi*, which entirely lacks it. Interestingly, the DH and NLS motifs of Cenp-C2 are more similar to those of *Sophophora* Cenp-C1 than to those of *Drosophila* Cenp-C1 (table 1). For the logo representation of the motifs, see Supplementary Figure S1.

**Table 1.** Genetic distances between the *Cenp-C* paralogs.

|  | *Sophophora* Cenp-C1 | *Drosophila* Cenp-C1 | *Drosophila* Cenp-C2 | Overall |
| --- | --- | --- | --- | --- |
| R-rich | 0.304 (0.254) | 0.435 (0.593) | 0.267 (0.244) | 0.463 (0.513) |
| DH | 0.292 (0.295) | 0.304 (0.380) | 0.283 (0.316) | 0.394 (0.445) |
| AT1 | 0.453 (0.505) | - | - | - |
| NLS | 0.281 (0.203) | 0.386 (0.443) | 0.284 (0.275) | 0.402 (0.413) |
| CenH3 | 0.316 (0.353) | 0.237 (0.232) | 0.254 (0.266) | 0.352 (0.371) |
| AT2 | 0.421 (0.530) | - | 0.301 (0.390) | 0.419 (0.498) |
| Cupin | 0.334 (0.402) | 0.294 (0.372) | 0.236 (0.283) | 0.422 (0.517) |
| Full-sequence | 0.404 (0.487) | 0.375 (0.484) | 0.363 (0.458) | 0.511 (0.634) |

Note – Values refer to distances between coding DNA sequences (values between brackets refer to amino acids distances).

**Figure 3.**
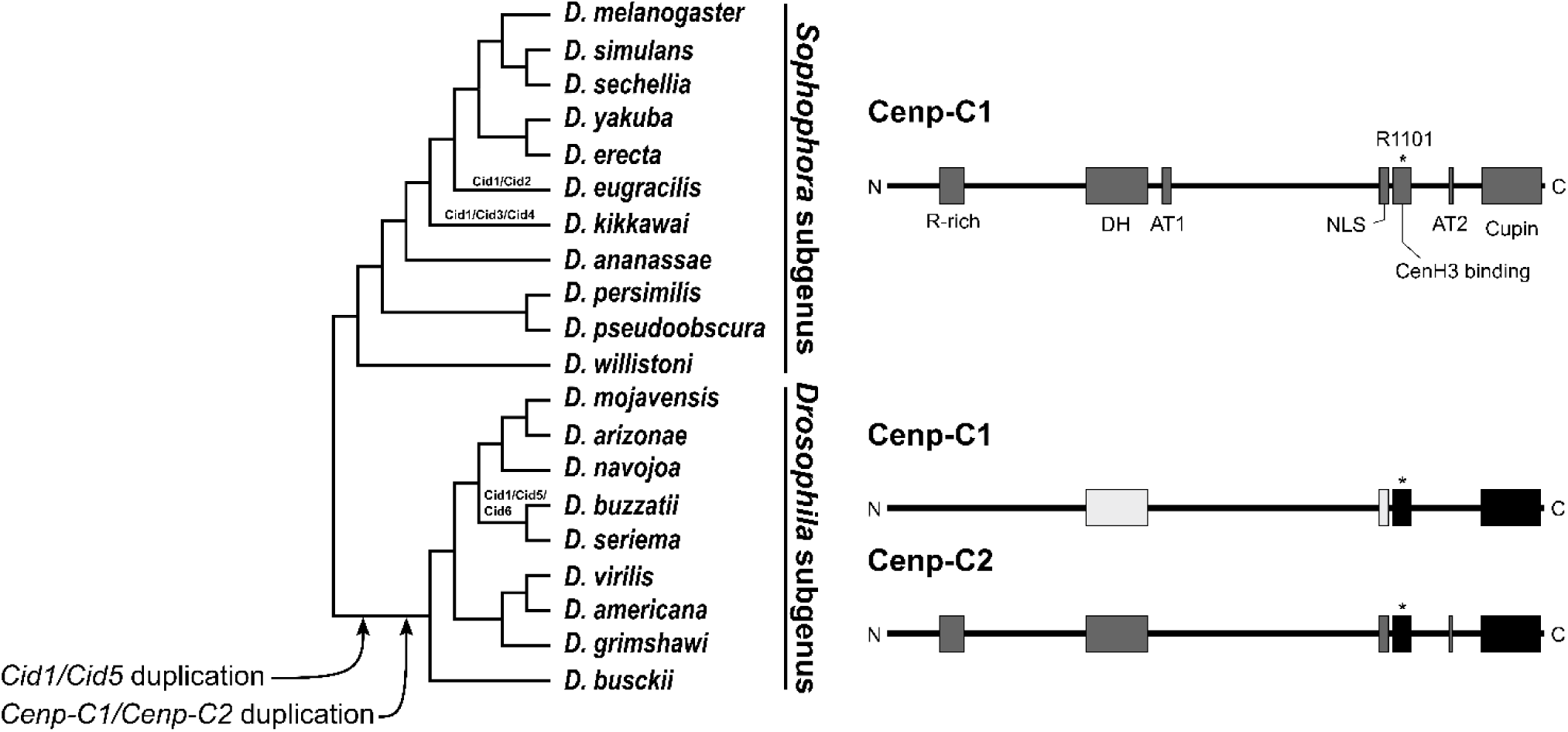
Some Cenp-C motifs are alternatively conserved between Cenp-C1 and Cenp-C2. Both *Cid* and *Cenp-C* genes were duplicated in the lineage that gave rise to species of the *Drosophila* subgenus, as indicated in the species tree. Moreover, *Cid1* was also duplicated in *D. eugracilis*, the *montium* subgroup (which includes *D. kikkawai*), and the *buzzatii* species cluster, the new paralogs of which are indicated at their respective branches. After the *Cenp-C* duplication, some functional motifs were alternatively conserved between the paralogs, as indicated at the right half of the image. High amino acids identity is indicated by the same color shade. Motifs are as follow: R-rich, arginine-rich; DH, drosophilid Cenp-C homology; AT1, AT hook 1; NLS, nuclear localization signal; CenH3 binding, also known as Cenp-C motif; AT2, AT hook 2; Cupin, a dimerization domain near the C-terminal region. The asterisk in the CenH3 binding motif indicates the corresponding R1101 of *D. melanogaster*, which is crucial for the centromere localization of Cenp-C1.

The conservation of the DH motif (involved in the recruitment of kinetochore proteins) and the NLS and CenH3 binding motifs (involved in centromere localization) in both Cenp-C1 and Cenp-C2 (fig. 3) indicates that it is unlikely that any of the paralogs underwent neofunctionalization. The (partial) loss of the Cupin motif in *D. busckii* and *D. grimshawi* points towards subfunctionalization. It is currently difficult to evaluate the loss of the AT1 motif in both Cenp-C1 and Cenp-C2, given that its function is unknown. However, the higher similarity of the DH and NLS motifs of Cenp-C2 to those of *Sophophora* Cenp-C1, the loss of the R-rich and AT2 motifs in Cenp-C1, and their retention in Cenp-C2 are highly indicative of subfunctionalization.

### The *Cenp-C* paralogs are differentially expressed

Given that Cenp-C is incorporated onto centromeres concomitantly with Cid (Schuh *et al*. 2007) and that the excess of both proteins can cause centromere expansion and kinetochore failure (Schittenhelm *et al*. 2010), the expression of both proteins needs to be tightly regulated. Kursel and Malik (2017) showed that *Cid5* expression is male germline-biased and proposed that *Cid1* and *Cid5* subfuncionalized and now performed nonredundant centromeric roles. In order to investigate if *Cenp-C1* and *Cenp-C2* are differentially expressed and correlated in some way with the expression of the *Cid* paralogs, we analyzed the available transcriptomes from embryos, larvae, pupae and adult females and males of *D. buzzatii* (Guillén *et al*. 2014), and from testes of *D. virilis* and *D. americana* (BioProject Accession PRJNA376405).

While *Cid6* is transcribed in all stages of development in *D. buzzatii*, confirming that *Cid6* functionally replaced *Cid1, Cid5* transcription is limited to pupae and adult males, with a higher transcription than *Cid6* in the latter (fig. 4A). Additionally, *Cid5* transcription is elevated in testes of *D. virilis* and *D. americana*, whereas *Cid1* is virtually silent (fig. 4C). Our results further support the finding of Kursel and Malik (2017) that *Cid5* displays a male germline-biased expression. In this context, our finding that *Cid5* is also transcribed in pupae of *D. buzzatii* may be related to the ongoing development of the male gonads.

**Figure 4.**
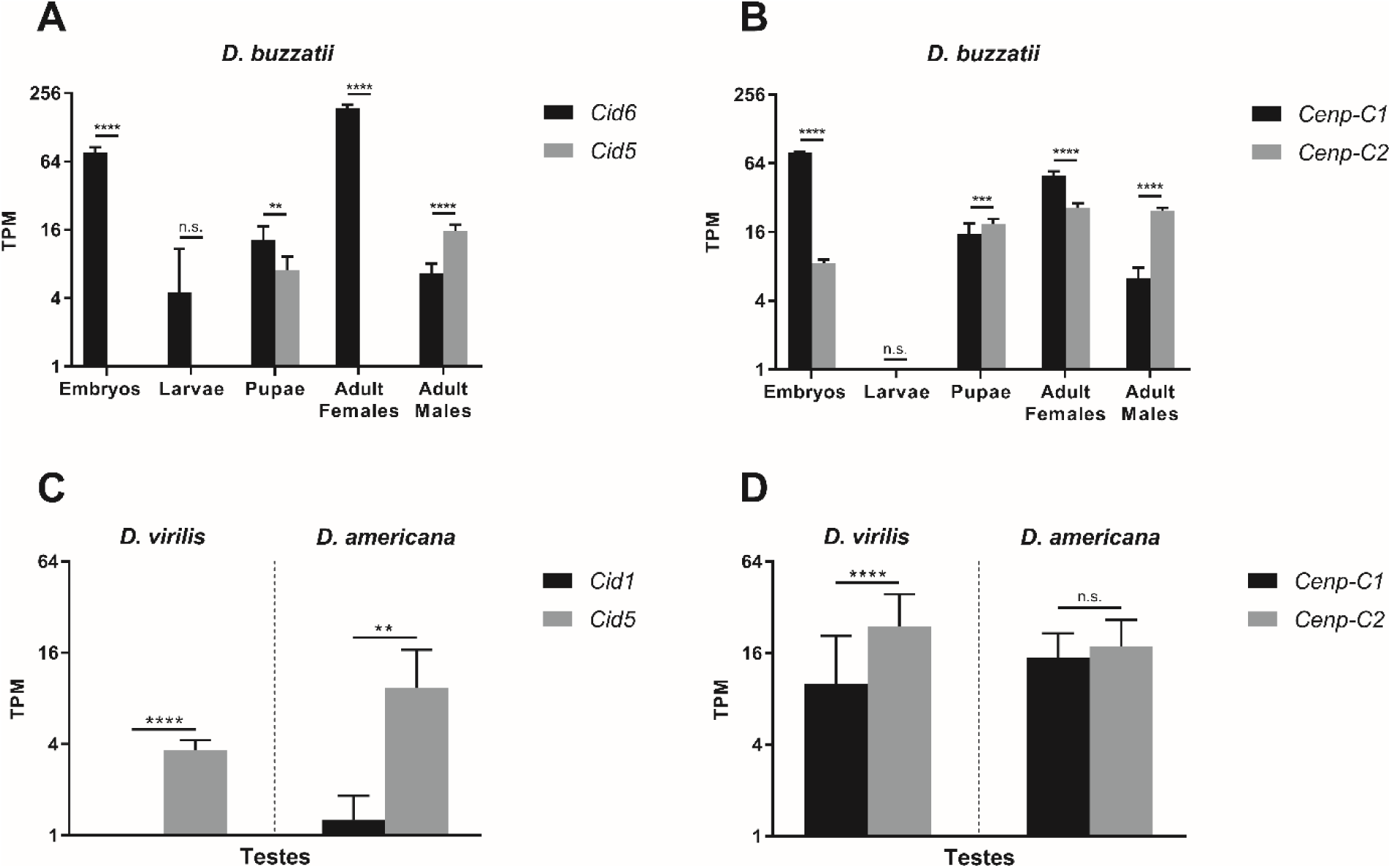
*Cid5* and *Cenp-C2* are male germline-biased. *Cid* and *Cenp-C* expression patterns in *D. buzzatii* (A and B) and *D. virilis* and *D. americana* (C and D). Data are presented as mean ± 95% confidence interval and analyzed by one-way ANOVA (A and B) and Student’s t-test (C and D): n.s., not significant; \**P* ≤ 0.05; \*\**P* ≤ 0.01; \*\*\**P* ≤ 0.001; \*\*\*\**P* ≤ 0.0001. TPM, transcripts per million.

In contrast to the *Cid* paralogs, we found that both *Cenp-C1* and *Cenp-C2* are transcribed in almost all stages of *D. buzzatii* development, with the exception of larvae (fig. 4B). *Cenp-C1* transcription is higher than that of *Cenp-C2* in *D. buzzatii* embryos and adult females. On the other hand, transcription of *Cenp-C2* is higher than that of *Cenp-C1* in *D. buzzatii* pupae and adult males. *Cenp-C2* transcription is also higher than that of *Cenp-C1* in *D. virilis* testes, but there is no significant difference between their expression in *D. americana* testes (fig. 4D). Therefore, similarly to the findings for the *Cid* paralogs, the differential expression between the *Cenp-C* paralogs in testis supports the subfunctionalization hypothesis. The male germline-biased expression of both *Cid5* and *Cenp-C2* points towards their interaction in spermatogenesis, but biochemical assays need to be performed to confirm this possible interdependence.

### The *Cid* and *Cenp-C* paralogs show signs of positive selection in species of the *repleta* group

The centromere drive hypothesis states that CenH3 and Cenp-C constantly evolve in an effort to suppress and diminish the associated deleterious effects of cenDNA selfish spread throughout the population by female meiotic drive (Henikoff *et al*. 2001; Dawe and Henikoff 2006). However, it has been proposed that the rapid evolution of CenH3 required for the “drive suppressor” function may be disadvantageous for canonical functions (e.g., mitosis; Finseth *et al*. 2015; Kursel and Malik 2017). The possibility that the paralogs achieved fitness optima for divergent functions predicts that selection may act differently in each of the *Cid* and *Cenp-C* paralogs. To test this hypothesis, we looked in our full-length alignments of the *Cid* and *Cenp-C* paralogs for signatures of positive selection using maximum likelihood methods. Given that CenH3 and Cenp-C are highly divergent, we focused our analyses on five closely related cactophilic *Drosophila* species from the *repleta* group (*D. mojavensis, D. arizonae, D. navojoa, D. buzzatii* and *D. seriema*).

We first used random-site and branch-site models to test for positive selection on particular sites during the evolution of the paralogs. The random-site models, which allow ω to vary among sites but not across lineages, revealed that both *Cid5* and *Cenp-C2* show extensive signs of positive selection (table 2). Particularly, Bayes Empirical Bayes analyses identified with a posterior probability > 95% four amino acids in the NTT of Cid5 and six amino acids across Cenp-C2 as having evolved under positive selection. Of the six Cenp-C2 amino acids, one is in the DH motif, one is in the Cupin motif, and the remaining four are in inter-motif sequences.

**Table 2.** Summary of random-site models for positive selection performed on each *Cid* and *Cenp-C* paralog.

|  | Alignment length (#nts) | M1a vs. M2a | M7 vs. M8 | M8a vs. M8 |
| --- | --- | --- | --- | --- |
| *Cid1/Cid6* | 609 | $P = 1$ | $P = 1$ | $P = 0.982$ |
| *Cid5* | 600 | $P = 0.099$ | $P = 0.069$ | **$P = 0.025$** |
| *Cenp-C1* | 3,492 | $P = 0.496$ | $P = 0.163$ | $P = 0.210$ |
| *Cenp-C2* | 3,696 | $P = 0.194$ | **$P = 0.005$** | $P = 0.068$ |

The branch-site models allow ω to vary both among sites and across branches on the tree and aim to detect positive selection affecting a few sites along particular lineages. The tests revealed that the paralogs show signs of positive selection in the branches of *D. navojoa Cid1* and *Cenp-C2, D. buzzatii Cenp-C1* and *Cenp-C2*, and *D. seriema Cenp-C1* and *Cenp-C2* (table 3). Particularly, Bayes Empirical Bayes analyses identified with posterior probability > 60% four amino acids in the NTT of *D. navojoa* Cid1, seven in inter-motif sequences of *D. navojoa* Cenp-C2, four in *D. buzzatii* Cenp-C1 (one in the DH motif and three in inter-motif sequences), six in inter-motif sequences of *D. buzzatii* Cenp-C2, four in *D. seriema* Cenp-C1 (two in the Cupin motif and two in inter-motif sequences), and six in inter-motif sequences of *D. seriema* Cenp-C2.

**Table 3.** Summary of branch-site models for positive selection performed on each *Cid* and *Cenp-C* paralog.

|  | MA vs. MAnull |  |  |  |  |  |
| --- | --- | --- | --- | --- | --- | --- |
|  | #1 | #2 | #3 | #4 | #5 | #6 |
| *Cid1* | $P = 1$ | **$P = 1.34\text{E-}06$** | $P = 1$ | $P = 1$ | $P = 0.251$ | $P = 1$ |
| *Cid5* | $P = 0.215$ | $P = 1$ | $P = 1$ | $P = 1$ | $P = 1$ | $P = 1$ |
| *Cenp-C1* | $P = 1$ | $P = 0.303$ | $P = 1$ | **$P = 0.0328$** | **$P = 1.08\text{E-}04$** | $P = 1$ |
| *Cenp-C2* | $P = 1$ | **$P = 1.64\text{E-}05$** | $P = 0.139$ | **$P = 0.041$** | **$P = 0.03$** | $P = 0.28$ |

Note – Foreground branches are as follow: #1 (*D. arizonae*, *D. mojavensis*); #2 (*D. navojoa*); #3 ((*D. arizonae*, *D. mojavensis*), *D. navojoa*); #4 (*D. buzzatii*); #5 (*D. seriema*); #6 (*D. buzzatii*, *D. seriema*).

Finally, we used clade model C to test for divergent selection among a priori designated lineages. The test reveal evidence of divergent selection acting on *Cid1, Cenp-C1* and *Cenp-C2* across almost all the foreground branches, the exception being *D. buzzatii* (Table 4). It is clear that the majority of sites are under negative selection across all lineages, and a small proportion do show signatures of positive selection (data not show); however, there is no obvious pattern of divergent selection across the phylogeny. Unlike the sites-models, clade models freely estimate ω‘s for each a priori designated clade and permit sites under positive selection in null models, which could explain the discrepancy among the sites-models and the clade model. Overall, we interpret our data as providing strong support for adaptive evolution at several sites in both the *Cid* and *Cenp-C* paralogs.

**Table 4.** Summary of the clade model for divergent selection performed on each *Cid* and *Cenp-C* paralog.

|  | CmC vs. M2a_rel |  |  |  |  |  |
| --- | --- | --- | --- | --- | --- | --- |
|  | #1 | #2 | #3 | #4 | #5 | #6 |
| *Cid1* | $P = 0.0575$ | $P = 0.048$ | **$P = 0.016$** | $P = 0.129$ | **$P = 0.009$** | $P = 0.022$ |
| *Cid5* | $P = 0.180$ | $P = 0.536$ | $P = 0.309$ | $P = 0.159$ | $P = 0.918$ | $P = 0.498$ |
| *Cenp-C1* | **$P = 0.0006$** | $P = 0.363$ | **$P = 0.039$** | $P = 0.072$ | $P = 0.108$ | **$P = 0.044$** |
| *Cenp-C2* | **$P = 0.00005$** | $P = 0.005$ | $P = 0.068$ | $P = 0.227$ | **$P = 0.011$** | $P = 1$ |

Note – Foreground branches are as follow: #1 (*D. arizonae*, *D. mojavensis*); #2 (*D. navojoa*); #3 ((*D. arizonae*, *D. mojavensis*), *D. navojoa*); #4 (*D. buzzatii*); #5 (*D. seriema*); #6 (*D. buzzatii*, *D. seriema*).

Our tests revealed that both the *Cid* and *Cenp-C* paralogs show signs of positive selection to some extent. Random-site models revealed that, on average, *Cid5* and *Cenp-C2* show extensive signs of positive selection, which may indicate that these male germline-biased genes possess drive-suppression function. Kursel and Malik (2017) found signs of positive selection in the *Cid3* paralog of the *montium* subgroup and proposed that *Cid3* and *Cid5* could be attenuating deleterious effects of centromere drive due to their male germline-biased expression. Our results of extensive positive selection on both *Cid5* and *Cenp-C2* do support this hypothesis. However, male germline-biased genes are widely known to evolve adaptively as the result of male-male or male-female competition (Ellegren and Parsch 2007; Meisel 2011). On the other hand, branch-site models revealed that different sites of both *Cenp-C1* and *Cenp-C2* show signs of positive selection in *D. buzzatii* and *D.seriema*, which may indicate that drive-suppression functions are not restricted to male-biased genes. Either way, molecular genetic data alone cannot reveal the underlying cause of adaptive evolution. What our findings do suggest is that species of the *Drosophila* subgenus likely have a specific inner kinetochore composition that mainly functions in spermatogenesis.

## Concluding remarks

The extensive diversity of kinetochore compositions in eukaryotes poses numerous questions regarding the flexibility of essential cellular functions (van Hooff *et al*. 2017). Is the kinetochore less conserved than other core eukaryotic cellular systems? And if so, why so many core kinetochore proteins are so diverse? Are the variants adaptive to the species? To answer such questions, it is necessary to investigate how a specific kinetochore composition affects specific cellular features and lifestyles. Herein, we showed that Cid5 and Cenp-C2 offer such a possibility, as both are inner kinetochore protein variants likely specialized to function mainly in spermatogenesis. Thus, finding out if and how Cid5 and Cenp-C2 play a role either in centromere drive suppression or reproductive competition can shed a new light into our understanding of centromere evolution.

## Materials and methods

### Identification of *Cid* and *Cenp-C* orthologs and paralogs in sequenced genomes

For most *Drosophila* species, *Cid* and *Cenp-C* coding sequences were obtained from EST data. For *Cenp-C1* of *D. navojoa, D. mojavensis, D. buzzatii, D. seriema* and *D. americana, Cenp-C2* of *D. buzzatii, D. seriema, D. americana* and *D. grimshawi, Cid5* of *D. virilis*, and both *Cid5* and *Cid6* of *D. buzzatii* and *D. seriema*, coding sequences were identified by tBLASTx in sequenced genomes. Since Cid is encoded by a single exon in *Drosophila*, we selected the entire open reading frame for each *Cid* gene hit, and since *Cenp-C* has multiple introns, we used the Augustus gene prediction algorithm (Stanke and Morgenstern 2005) to identify the coding DNA sequences. For annotated genomes, we recorded the 5’ and 3’ flanking genes for the *Cid* and *Cenp-C* genes of each species. For genomes that are not annotated, we used the 5’ and 3’ nucleotide sequences flanking the *Cid* and *Cenp-C* genes as queries to the *D. melanogaster* genome using BLASTn and verified the synteny in accordance to the hits. For the *D. seriema* genome assembly, see Supplementary File S1. All *Cid* and *Cenp-C* coding sequences and their database IDs can be found in Supplementary Files S2 and S3, respectively.

### Fluorescent *in situ* hybridizations (FISH) on polytene chromosomes

Probes for *Cid1* /*Cid6* were obtained by PCR (see fig. 1A for primer site) from genomic DNA of *D. buzzatii* (strain st-1), *D. seriema* (strain D73C3B), *D. mojavensis* (strain 14021-0248.25) and *D. virilis* (strain 15010-1551.51). We cloned the PCR products into the pGEM-T vector (Promega) and sequenced them to confirm identity. Recombinant plasmids were labeled with digoxigenin 11-dUTP by nick translation (Roche Applied Science). FISH on polytene chromosomes was performed as described in Dias *et al*. (2015). The slides were analyzed under an Axio Imager A2 epifluorescence microscope equipped with the AxioCam MRm camera (Zeiss). Images were captured with the AxioVision software (Zeiss) and edited in Adobe Photoshop. Chromosome arms were identified by their morphology (Kuhn *et al*. 1996; González *et al*. 2005; Schaeffer *et al*. 2008).

### Phylogenetic analyses

*Cid* and *Cenp-C* sequences were aligned at the codon level using MUSCLE (Edgar 2004) and refined manually. Subsequently, we generated maximum likelihood phylogenetic trees in MEGA6 (Tamura *et al*. 2013) with the GTR substitution model and 1,000 bootstrap replicates for statistical support.

### Expression analyses

RNA-seq data from *D. buzzatii* (Guillén *et al*. 2014), and from *D. virilis* and *D. americana* (BioProject Accession PRJNA376405) were analyzed for the *Cid* and *Cenp-C* expression patterns with Bowtie2 (Langmead and Salzberg 2012), as implemented to the Galaxy server (Afgan *et al*. 2016). Mapped reads were normalized by the transcripts per million (TPM) method (Wagner *et al*. 2012), and all normalized values < 1 were set to 1 so that log2 TPM ≥ 0.

### Positive selection analyses

*Cid* and *Cenp-C* alignments and gene trees were used as input into the CodeML NSsites models of PAMLX version 1.3.1 (Xu and Yang 2013). Random-site and branch-site models were used to test for positive selection on particular sites during the evolution of the *Cid* and *Cenp-C* paralogs. Random-site models allow ω to vary among sites but not across lineages; for this analysis, we compared three models that do not allow ω to exceed 1 (M1a, M7 and M8a) to two models that allow ω > 1 (M2a and M8). Branch-site Model A was compared with Model A_null_ to examine whether particular sites evolved under positive selection along a priori specified branches (called foreground branches). Foreground branches were as follow: #1 (*D. arizonae, D. mojavensis*); #2 (*D. navojoa*); #3 ((*D. arizonae, D. mojavensis*), *D. navojoa*); #4 (*D. buzzatii*); #5 (*D. seriema*); #6 (*D. buzzatii, D. seriema*). Positively selected sites were classified as those with a Bayes Empirical Bayes posterior probability > 90%. Clade model C (CmC) tests for divergent selection on particular sites among a priori designated lineages. The modified null model of CmC (M2a_rel) assumes that sites fall into three classes: purifying selection (0 < ω < 1); neutral evolution (ω = 1); or positive selection (ω >1). In CmC, the third site class allows the estimated ω for a site to diverge across foreground branches. Foreground branches were as follow: #1 ((*D. arizonae, D. mojavensis*), *D. navojoa*); #2 (*D. buzzatii, D. seriema*).

## Acknowledgments

We are grateful to Dr. Maura Helena Manfrin (Univesity of São Paulo) for providing us the *D. seriema* strain. This work was supported by a grant from “Fundação de Amparo à Pesquisa do Estado de Minas Gerais” (FAPEMIG) to G.K. (grant number APQ-01563-14).

---

**Supplementary Figure S1.**
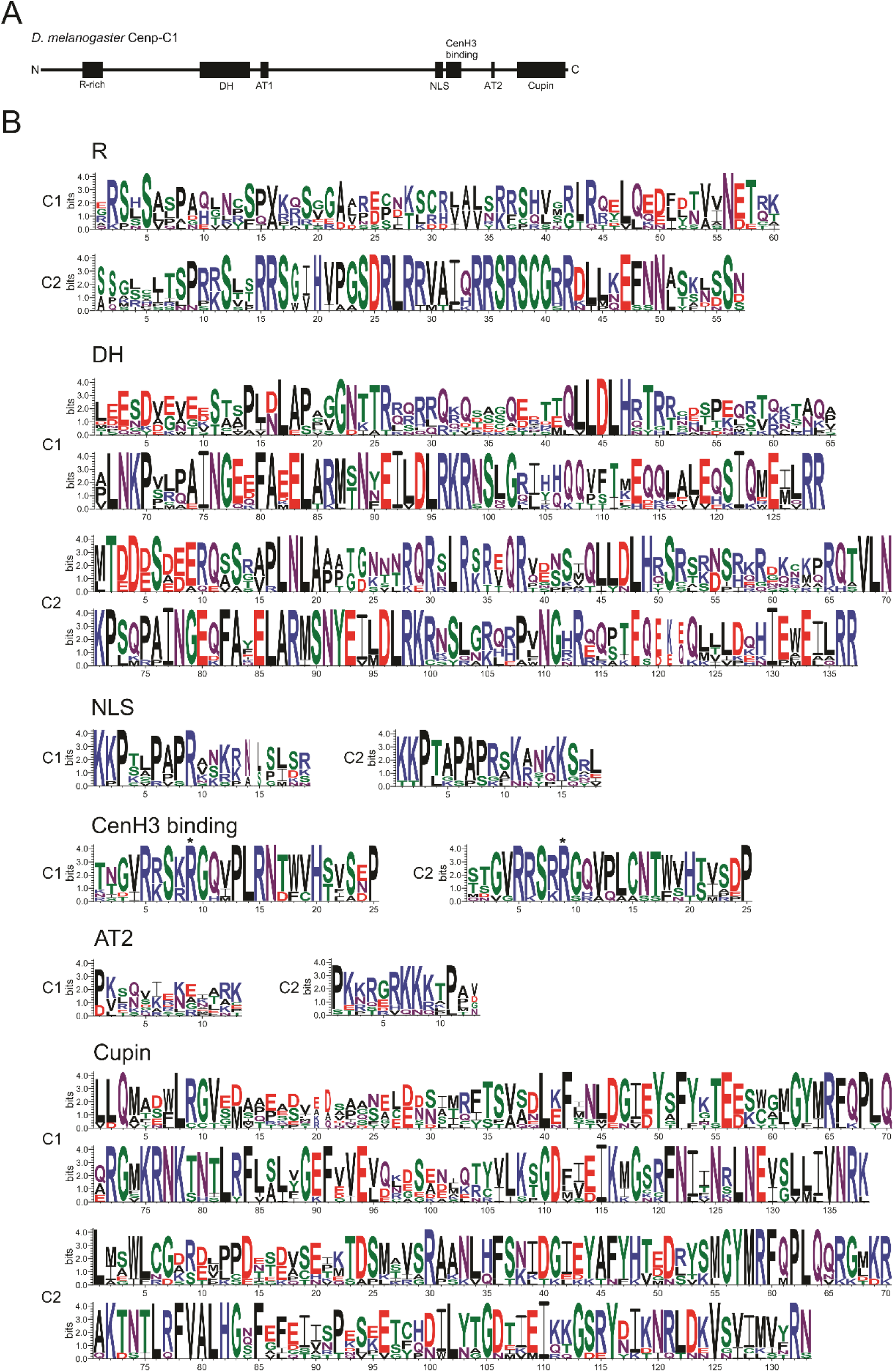
Some Cenp-C motifs are alternatively conserved between Cenp-C1 and Cenp-C2. (A) Schematic representation of the motif structure of D. melanogaster Cenp-C1. (B) Logo representations for each motif of the Drosophila subgenus Cenp-C1 (C1) and Cenp-C2 (C2). Motifs are as follow: R-rich, arginine-rich; DH, drosophilid Cenp-C homology; AT1, AT hook 1; NLS, nuclear localization signal; CenH3 binding, also known as Cenp-C motif; AT2, AT hook 2; Cupin, a dimerization domain near the C-terminal region. The asterisk in the CenH3 binding motif indicates the corresponding R1101 of D. melanogaster, which is crucial for the centromere localization of Cenp-C1.

